# A cold-blooded vertebrate shows integration of antimicrobial defenses and tissue repair through fever

**DOI:** 10.1101/2022.10.27.513985

**Authors:** Farah Haddad, Amro M. Soliman, Michael E. Wong, Emilie H. Albers, Shawna L. Semple, Débora Torrealba, Ryan D. Heimroth, Asif Nashiry, Keith B. Tierney, Daniel R. Barreda

**Author notes:** These authors contributed equally to this work (Farah Haddad, Amro M. Soliman). For correspondence: Daniel R. Barreda, Department of Biological Sciences, University of Alberta, Canada.

## Abstract

Multiple lines of evidence support the value of moderate fever to host survival, but the mechanisms involved remain unclear. This is difficult to establish in warm-blooded animal models, given the strict programs controlling core body temperature and the physiological stress that results from their disruption. Thus, we took advantage of a cold-blooded teleost fish that offered natural kinetics for the induction and regulation of fever and a broad range of tolerated temperatures. A custom swim chamber, coupled to high-fidelity quantitative positional tracking, showed remarkable consistency in fish behaviours and defined the febrile window. Animals exerting fever engaged pyrogenic cytokine gene programs in the CNS, increased efficiency of leukocyte recruitment into the immune challenge site, and markedly improved pathogen clearance *in vivo*, even when an infecting bacterium grew better at higher temperatures. Contrary to earlier speculations for global upregulation of immunity, we identified selectivity in the protective immune mechanisms activated through fever. Fever then inhibited inflammation and markedly improved wound repair. Artificial mechanical hyperthermia, often used as a model of fever, recapitulated some but not all benefits achieved through natural host-driven dynamic thermoregulation. Together, our results define fever as an integrative host response that regulates induction and resolution of acute inflammation, and demonstrate that this integrative strategy emerged prior to endothermy during evolution.

## INTRODUCTION

Fever is a cornerstone of acute inflammation. ^1^ The classical response is initiated through the recognition of damage-associated molecular patterns (DAMPs) or pathogen-associated molecular patterns (PAMPs) by pattern recognition receptors (PRRs) on the surface of immune leukocytes. This sensing reaction leads to the activation of resident myeloid cells at the challenge site, and is followed by rapid production of pyrogenic prostaglandin E2 (PGE2) and cytokines, including tumor necrosis factor-alpha (TNF-α), interleukin-1 beta (IL-1β), and interleukin-6 (IL-6). ^2^ Contributions at multiple levels of the fever cascade are observed; IL-6, for example, promotes initial activation events, the early rise in core body temperature, as well as the subsequent orchestration of lymphocyte trafficking to lymphoid organs. ^3^ Others, like interleukin-8 (IL-8) are produced locally at the site of infection and manage recruitment of inflammatory leukocytes from circulation, peripheral stores and the hematopoietic compartment. ^4^ Increased synthesis of cyclooxygenase 2 (COX-2) within the median preoptic nucleus of the hypothalamus fosters added production of PGE2, which serves a dominant pyrogenic role in fever. ^5^ Production of endogenous pyrogenic cytokines such as IL-1β and IL-6 within the central nervous system (CNS) may also complement the activity of those generated in peripheral tissues during the induction of fever. ^3^ Systemic physiological changes in vasodilation, vascular permeability and leukocyte recruitment become evident a few hours after the initial insult. ^6^ Thus, early innate immune recognition initiates a highly organized response that engages neuronal circuitry in the central and peripheral nervous systems and triggers the activation of thermoregulatory pathways.

The association of fever and disease dates back at least as far as Hippocrates (2500 years). ^7^ A rise in body temperature is so tightly associated with the inflammatory response that heat (*calor*) is one of the four cardinal signs of inflammation. While non-severe forms of fever are well established to increase host survival upon infection, ^3, 8, 9^ the mechanisms behind these contributions remain poorly understood. Improved host protection has been postulated to stem from a direct impact of increased temperature on invading pathogens, and global upregulation of antimicrobial immune mechanisms. ^3, 10^ For thermal restriction, pathogen survival and replication can be directly compromised when their maximum tolerated temperatures are reached or exceeded. ^10^ This is well documented for many microbes and served as an effective therapy against diseases like neurosyphilis and gonorrhea prior to the advent of antibiotics. ^9^ However, many pathogens are known to be largely unaffected or grow better at the higher temperatures that fever elicits. ^10, 11^ And, even for originally permissive organisms, the effectiveness of thermal restriction may be limited to initial encounters between pathogen and host, given the extensive repertoire of thermal resistance mechanisms that are available to viruses, archaea, bacteria, fungi, and parasites. ^10, 11^ Global promotion of immune defenses has also been proposed, based on reported effects of temperature increases on metabolic rates and multiple effectors and regulators of innate and adaptive immunity. ^3, 12^ However, the global nature of this induction contradicts the emphasis that a host is known to place on energy conservation and management of collateral inflammation-associated tissue damage. ^13, 14^ As a result, debates on the net value of fever to host health continue to permeate the literature. ^7, 15–18^ This is compounded by limitations among available experimental models to adequately recapitulate the natural physiological processes driving and sustaining febrile responses. Early assessments into the benefits of fever, for example, suffered from temporal deviations - fever was artificially induced prior to infection. ^9, 12^ In other cases, thermal ranges outside those normally elicited by fever were used, or peak temperatures were maintained for extended periods of time. ^9, 12^ *In vitro* and *in vivo* mammalian models of fever-range hyperthermia (FRH) continue to offer valuable insights and have confirmed improvements in host survival and decreased microbial loads because of increased core body temperatures. ^3, 19^ Unfortunately, exogenous mechanical temperature manipulation is also well established to cause physiological stress and fails to replicate the host’s intrinsic thermoregulatory machinery for heating and cooling elicited during natural fever. ^16^ Pharmacological models based on antipyretic drug administration (e.g., NSAIDs) have also been used broadly, but are hampered by inhibition of inflammatory pathways at multiple points and other off-target effects. ^16, 20^ As a result, fever remains among the most poorly understood of the acute inflammatory processes.

Ectotherms (fish, amphibians, reptiles, invertebrates) and endotherms (mammals, birds) induce fever upon infection, and both exhibit a strong behavioural component. ^21, 22^ Common biochemical pathways appear to drive fever across ectothermic and endothermic animals. ^3, 23^ Ectotherms, however, rely on behaviour to induce fever in the absence of the metabolic toolkit available to endotherms. ^3, 24, 25^ Upon infection, fish move to areas with warmer waters, while reptiles lay on sun-warmed terrestrial environments. ^24^ Social animals like honeybees go further, displaying behavioural thermoregulation at the group level, to achieve a communal increase in hive temperature in response to infection. ^26^ In all, the conservation of fever across phylogeny spans 550 million years of metazoan evolution. ^24^ The net result is a survival advantage ^3, 20^ which, based on the long-standing natural selection of the fever response, appears to heavily out-weight the reported metabolic costs, ^24, 27^ increased potential for predation, ^28^ and decrease in reproductive success. ^29^ This level of commitment mirrors that displayed by some pathogens to inhibit it. Herpesvirus, for example, has been recently shown to express soluble decoy TNF receptors during infection that delay behavioural fever and allow for increased viral replication. ^30^

In this study we used a cold-blooded teleost vertebrate model to gain additional insights into the immunobiology of fever. We examined febrile responses under host-driven dynamic thermoregulation, to more closely mimic natural conditions for heating and cooling. This avoided common caveats encountered with exogenous drugs, temporal deviations from native thermoregulatory programs, or forcing animals beyond thermal ranges normally elicited through fever. An *in vivo Aeromonas* cutaneous infection model was tailored to focus on the most common moderate self-resolving form of this natural biological process rather than severe pathological fever. Under the experimental conditions used, fever was transient and self-limiting, enabling us to interrogate its potential contributions during induction and resolution phases of acute inflammation. Together, our results demonstrate that fever is not a biproduct of acute inflammation but an important regulator of its induction and control. The functional attributes of fever in ectotherms stem from earlier and selective rather than stronger induction of innate antimicrobial programs against infection. These are further integrated with efficient control of inflammation and promotion of wound repair.

## RESULTS

### High-resolution motion tracking identifies a predictable fever program in an outbred population of fish

Conventional shuttle box approaches for examination of fish behaviour are well established to produce significant variability between individual animals, driven by differences in their preference for cover, swim depth and activity level, in addition to social behaviours associated with schooling, territorial grouping or avoidance based on dominance. ^31^ In an effort to decrease this heterogeneity and offer greater analytical depth to our behavioural outputs, we customized an annular temperature preference tank (ATPT) ^32^ which takes advantage of fluid dynamics instead of physical barriers to establish distinct temperature environments (**Figure 1**). Under this setup, aquatic animals were free to choose environmental temperatures independent of other variables such as lighting, perceived cover, edge effects, water depth or current that impact positional behaviour. Distinct temperature set points (16, 19, 21, 23, 26°C) were chosen and used to create a barrier-free environmental housing temperature gradient that spanned a 10°C range (**Figure 1A-C**). Water inputs and flow rates were optimized to yield a gradient that was stable through 14 days of continuous evaluation (**Figure 1C**). Directional flow rates across the swim chamber were also adjusted to create smaller consistent temperature gradients between each of the primary thermal zones (**Figure 1B**). Our goal was to avoid abrupt housing water temperature boundaries, which could affect an animal’s choice to transition between adjacent thermal zones. Next, we coupled this updated ATPT to an automated monitoring system with per-second temporal resolution for effective tracking of fish through day and night cycles (**Figure 1D**). This offered greater analytical robustness and temporal resolution than previously possible in behavioural fever analyses. We then challenged individual teleost fish (*Carassius auratus*; goldfish) with an *Aeromonas* cutaneous infection *in vivo*. As eurythermal ectotherms, goldfish offered an opportunity to examine absolute changes to thermopreference in response to an immune challenge, while minimizing the potential for thermal stress. This is because the natural range of tolerated environmental temperatures for these fish (1.3°C to 34.5°C) was broader than those temperatures expected in a febrile response. Our choice also provided access to a previously optimized *in vivo* self-resolving animal model where changes during induction and resolution phases of acute inflammation could be examined. _33, 34_

**Figure 1.**
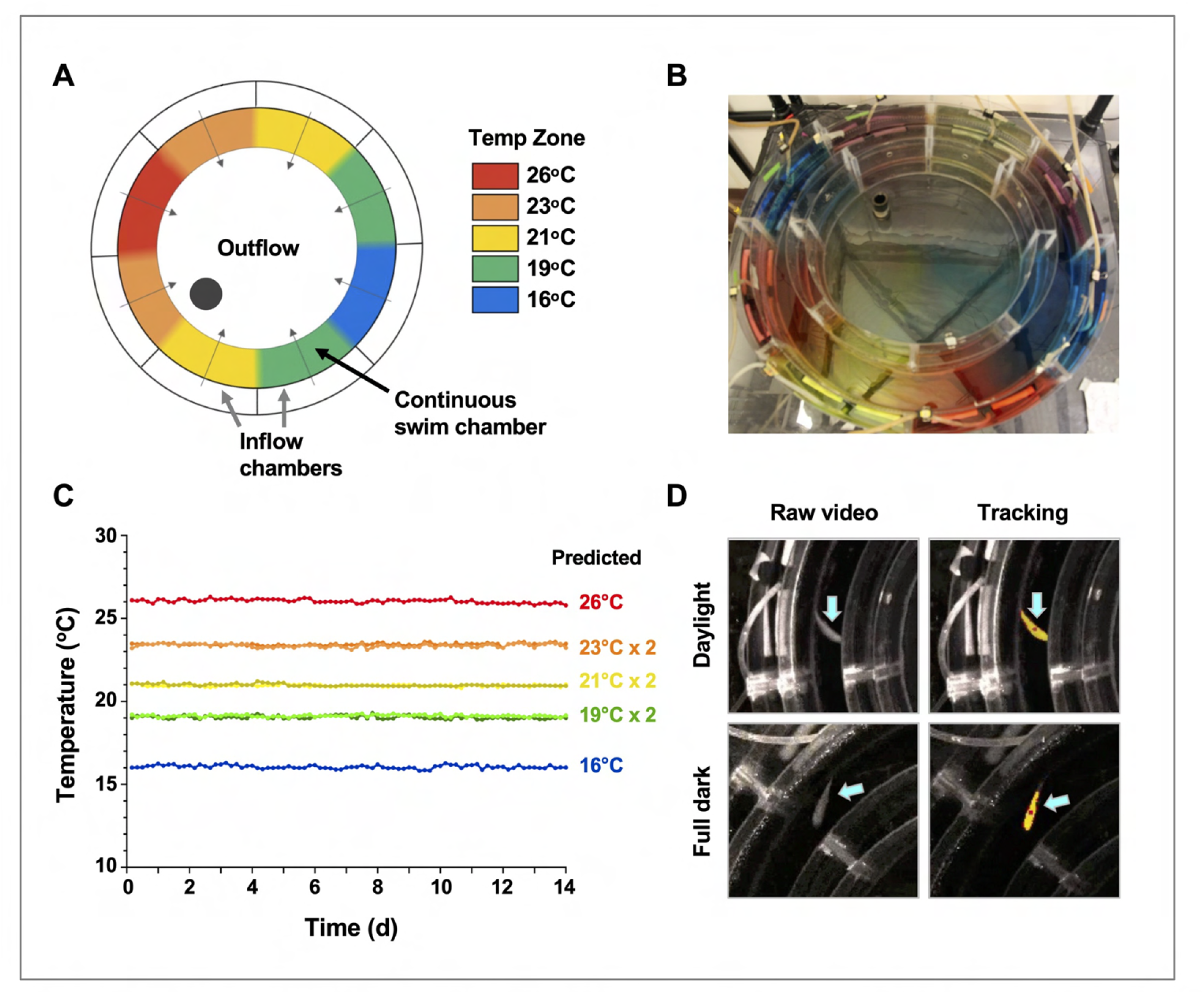
Annular temperature preference tank design, validation, and fish tracking. (A) The ATPT established a continuous ring-shaped swim chamber that offered distinct temperature environments separated by fluid dynamics instead of physical barriers. (B) Dye flow test highlights eight distinct thermal zones created using concentric flow directed towards the center of the apparatus. (C) Analysis of temperature stability for established thermal zones. Single lines correspond to highest (26°C) and lowest (16°C) temperatures. Double lines denote values from equivalent zones on opposing sides of the apparatus for 19°C, 21°C, and 23°C target temperatures. (D) Representative images of a fish (blue arrows) in raw infrared and processed video, under simulated daylight and night (full dark) conditions. Yellow identifier denotes strong tracking signal achieved for experimental setup. Red dot denotes center point used to set coordinates for raw behavioural data.

Behavioural examination identified four distinct phases of the fever response among groups of fish challenged *in vivo* with *Aeromonas veronii* (**Figure 2A**). Thermal selection patterns across challenged fish were remarkably reproducible, even when fish were placed individually in the annular swim chamber (**Figure 2B**). A distinct period from 1 to 8 days post-infection (dpi) emerged, where *Aeromonas* challenged fish displayed a 2-3°C increase in thermal preference compared to mock-infected (saline) controls (**Figure 2B**). Variance analysis confirmed the consistency in environmental temperature selected by individual animals within this 1-8 dpi time window (**Figure 2B**). Outside of this window, we found no significant difference in thermal selection between challenged and control individuals, with both *Aeromonas* and saline-treated groups also shifting to large fluctuations both temporally and among individual fish (**Figure 2B**).

**Figure 2.**
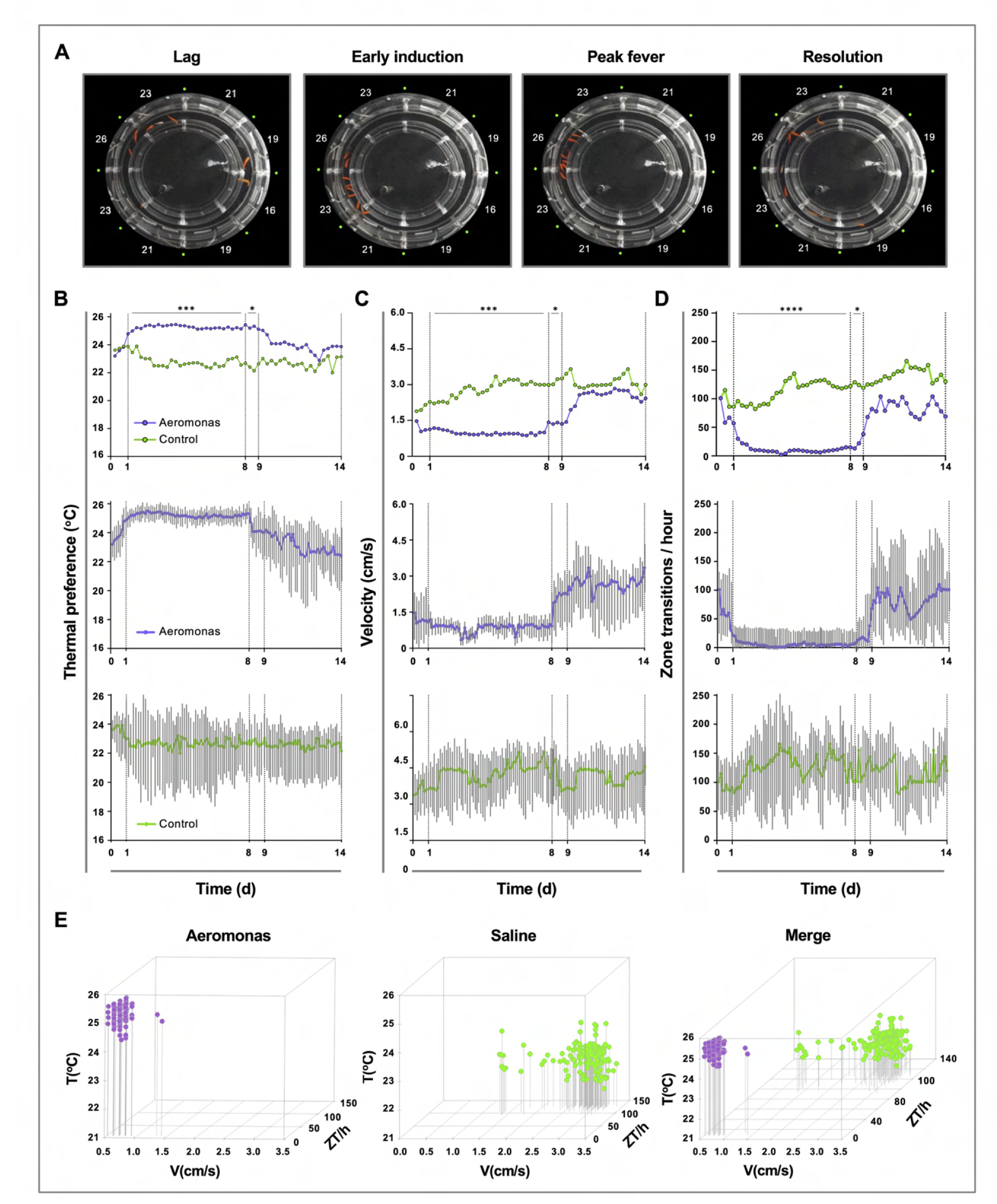
Homogeneity in thermal preference and sickness behaviours among fish eliciting fever. **(A)** Fish infected with *Aeromonas veronii* were free to select a range of environmental temperatures within the ATPT. Video still images show collective positioning of fish during distinct phases of fever response. Labels correspond to mean temperature for each ATPT thermal zone. **(B)** Temperature preference, **(C)** velocity and **(D)** total transitions across thermal zones for fish infected with *Aeromonas* (n=5) or mock-infected with saline (n=5). Fish were placed separately in the ATPT and individually monitored for 2 weeks. Evaluation of behavioural variance shows distinct periods of marked consistency in temperature preference, swimming velocity, and thermal zone transitions across *Aeromonas*-infected fish. Solid lines represent mean hourly values and vertical grey bars correspond to standard deviation at each time point. Results were analyzed by an ordinary two-way ANOVA and Šídák’s multiple comparisons test. \**p<*0.05, \*\**p<*0.01, \*\*\**p<*0.001, \*\*\*\**p<*0.0001). **(E)** Simultaneous three-parameter representation of behaviour hourly data points for *Aeromonas*-infected and saline control fish (n=5 fish and 168 data points per group; mean values for 5 fish shown). 3D plots correspond to 1-8 dpi febrile period. Merged graph highlights segregation of behavioural responses. Figure was produced using R (version 3.3); open-source code provided as supplementary material.

Our behaviour analyses further identified two new measurable lethargy-associated outcomes in teleost fish, which add to the similarities between ectotherm and endotherm fever. The first was defined by a decrease in swimming velocity (V) among *Aeromonas*-challenged fish (1-8 dpi; **Figure 2C**). In contrast, velocity among control fish remained high and variable across individuals during the same timeframe (**Figure 2C**). The second lethargy parameter was based on changes to temperature seeking behaviour, defined by the rate of transitions that a fish made between distinct ATPT thermal zones. Whereas control saline injected fish continued to show one hundred or more zone transitions (ZT) per hour, *Aeromonas*-treated fish displayed a dramatic decrease in the number of ZT during the same 1-8 dpi window (**Figure 2D**). As with temperature preference and velocity measurements, ZT values within this fever behavioural window were remarkably consistent across individual *Aeromonas*-challenged fish, increasing in variance after 8 dpi (**Figure 2D**). In sharp contrast, control fish displayed significant heterogeneity throughout the entire observation period. These two new lethargy-associated metrics are consistent with established sickness behaviours of metabolic fever in humans and other endotherms (immobility, fatigue, and malaise) and help to further define the behavioural fever response of teleost fish.

Hourly values for *Aeromonas*-challenged and saline control groups were evaluated simultaneously during the established fever window (1-8 dpi) and across the broader 14-day observation period. Between 1-8 dpi, we identified marked segregation in the responses elicited by fish in these two groups (**Figure 2E**). For *Aeromonas*-challenged fish, V and ZT values remained exclusively low during the febrile period (**Figure 2E**). In contrast, saline control fish exhibited a wider range of movement profiles, dominated by high V and ZT values (**Figure 2E**).

### Activation of CNS and systemic febrile programs following *Aeromonas* infection

To confirm classic fever engagement of the central nervous system via fever and assess potential differences with mechanical fever-range hyperthermia, hypothalamic tissue was isolated from *Aeromonas*-challenged fish and examined for local expression of pyrogenic cytokines. The selected genes, IL-1β, TNFα and IL-6, showed more robust local induction of gene expression under dynamic fever conditions (T_D_ group), where fish had been allowed to swim freely through the established 10°C temperature gradient within the ATPT (**Figure 3A**). Two cytoprotective elements (HSP70 and HSP90) further displayed the highest levels of expression in the hypothalamus under these dynamic thermal conditions. Evaluation of PGE_2_ concentrations in systemic circulation showed an early peak at 24 hours post-infection (hpi) for T_D_ fish, consistent with its role as a major pyrogenic mediator of fever ^3^ (**Figure 3B**). These responses were distinct from those fish placed under 26°C (T_S26_; mechanical fever-range hyperthermia) or 16°C static thermal conditions (T_S16;_ basal acclimated temperature) following infection. T_S26_ FRH promoted upregulation of cytokine and cytoprotective genes in our panel, but to lower levels than those fish allowed to exert dynamic fever. Circulating PGE_2_ concentrations remained near basal levels for both T_S26_ FRH and T_S16_ groups. Thermal increases alone (without pathogen stimulus) were not sufficient to activate these cytokines and cytoprotective genes in the CNS (**Figure S1**).

**Figure 3.**
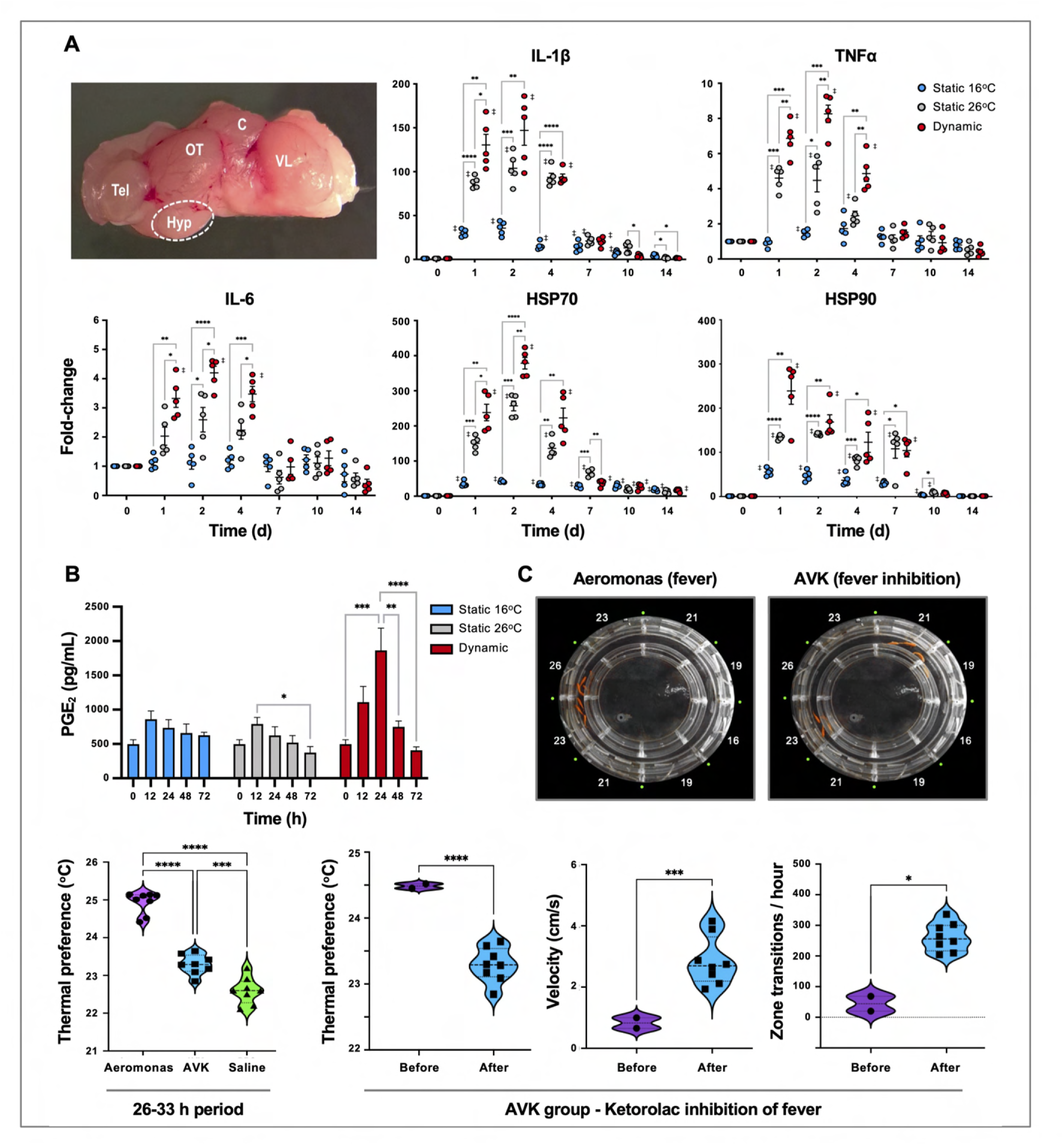
Confirmation that fish fever engages CNS and systemic pyrogenic signals following an *Aeromonas* infection. Fish were inoculated with *Aeromonas* and placed in static 16°C (basal acclimated temperature), static 26°C (mechanical hyperthermia; maximum temp. that fish selected during behavioural fever), or dynamic fever (where fish could move freely between thermal zones). **(A)** qPCR evaluation of hypothalamic responses following infection (n=5 per group per time point; 3 technical replicates per fish per time point). Symbols correspond to individual samples; lines represent the mean ± SEM. Results analyzed by an ordinary two-way ANOVA using Tukey post-hoc test. **(B)** Plasma collected at 0, 12, 24, 48 and 72 h post-infection (n=6 per group per time point) and PGE_2_ concentrations determined via ELISA. Results analyzed by an ordinary one-way ANOVA followed by a Tukey’s post-hoc test. Error bars represent +SEM. **(C)** Confirmation that an antipyretic inhibits fish fever. Video still images illustrate disruption of high thermal preference collective positioning following treatment of *Aeromonas*-infected fish with ketorolac tromethamine. Quantitation of thermal preference shows differences between fish infected with *Aeromonas*, those infected with *Aeromonas* and treated with ketorolac (AVK), and those mock-infected with saline (n=5 fish per group per time point). Observation period began 2 h after start of peak febrile response (26 h) and continued for 8 h duration of ketorolac action. One way ANOVA followed by Tukey’s multiple comparisons test compared *Aeromonas*, AVK, and saline groups. Changes were further evaluated within AVK group before and after ketorolac administration. Welch’s t-tests compared behavioural parameters. For all, \**p<*0.05, \*\**p<*0.01, \*\*\**p<*0.001, \*\*\*\**p<*0.0001; ‡ denotes significant difference from time 0, *p<*0.05.

Administration of an antipyretic offered added support for the shared biochemical pathways directing fever in ectotherms and endotherms. We chose ketorolac as a nonsteroidal anti-inflammatory drug (NSAID) with the capacity to inhibit COX-1 and COX-2. ^35^ This drug has been successfully used in a range of animal species ^36^ and can be injected, ^35, 37^ thereby ensuring consistent dosing. In humans, a dose of 0.5 mg/kg is effective, and results in a 15-20 min onset with a 6-8 h duration of action. ^35^ Similarly, injection of ketorolac to *Aeromonas*-infected fish inhibited fever at a dose of 0.5 mg/kg (Aeromonas versus AVK experimental group; **Figure 3C**). Examination of changes within the AVK group before and after ketorolac administration further showed that this NSAID inhibited fever-associated increases in thermal preference and lethargy behaviours (**Figure 3C**).

### Fever markedly improves *Aeromonas* clearance while showing selectivity in the induction of ROS and NO antimicrobial defenses

During the course of infection, furuncles caused by *Aeromonas* species can shed up to 10^7^ bacteria per hour in fish. ^38^ Thus, we assessed the presence of *A. veronii* on the furuncle surface as a measure of pathogen load and shedding potential. Infected fish held at 16°C static thermal conditions (T_S16_) displayed heavy bacterial loads through the first four days after infection, reducing to 70 ± 23 and 28 ± 10 CFU at 7 and 10 dpi, respectively, and subsequently progressing below detectable levels by 14 dpi (**Figure 4A**). Fish in the dynamic fever group (T_D_) also showed heavy initial bacterial burden, but these decreased markedly faster compared to T_S16_ fish (40 ± 27 CFU at 4 dpi, and below detectable levels by 7 dpi). Mechanical hyperthermia (T_S26_) yielded an intermediate response, achieving *Aeromonas* clearance 10 days following infection (**Figure 4A**). Thus, fish allowed to exert dynamic fever cleared *A. veronii* in half the time than fish maintained in static 16°C conditions. Notably, this enhancement in clearance could not be explained by the current thermal restriction model since *A. veronii* showed faster growth as incubation temperatures increased from 16 to 26°C (**Figure 4B**).

**Figure 4.**
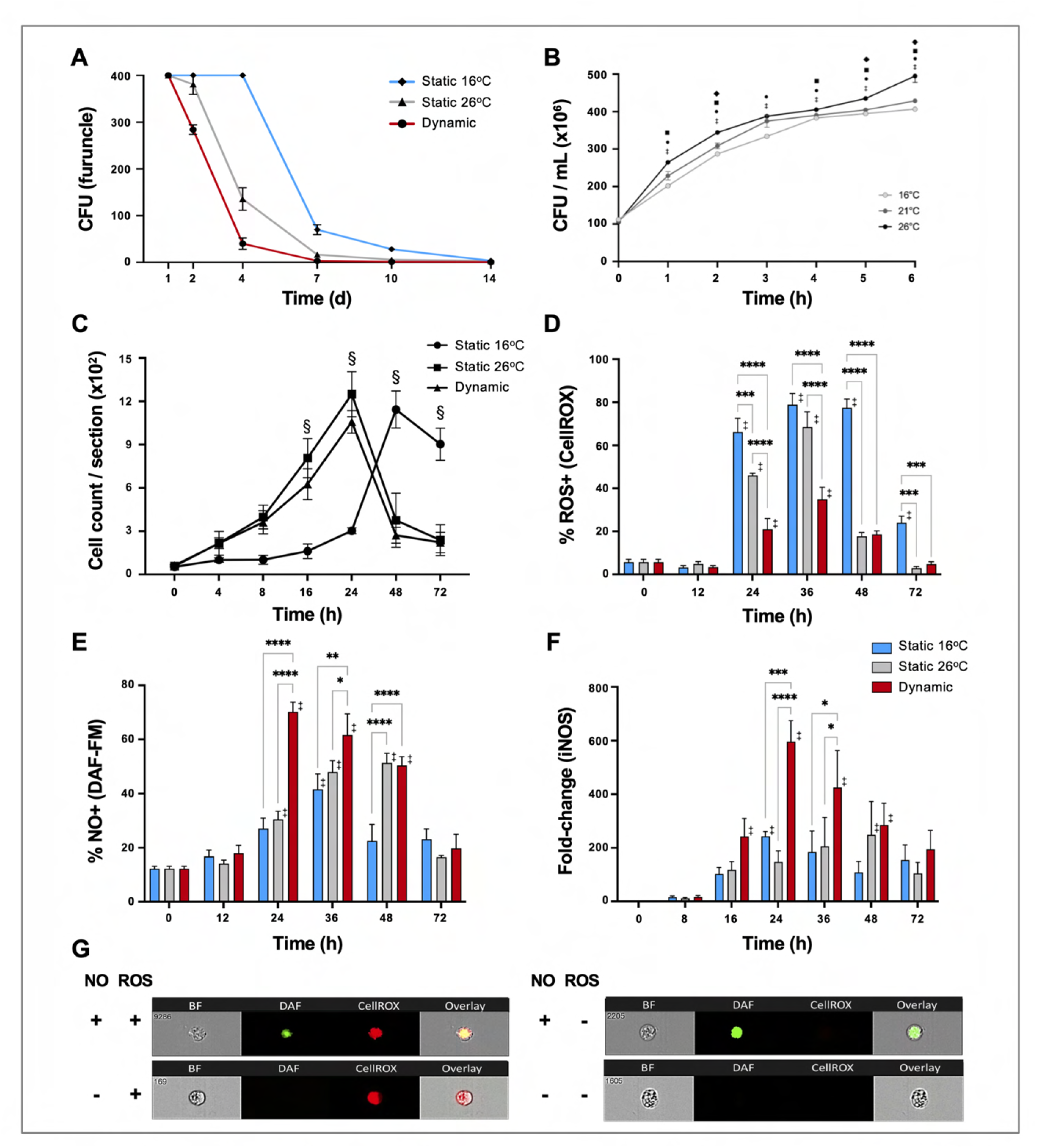
Fever enhances pathogen clearance while showing selectivity in the induction of ROS and NO antimicrobial defenses. Fish were infected with *Aeromonas* and placed under 16°C static, 26°C static (mechanical hyperthermia), or dynamic host-driven fever conditions. **(A)** Bacterial loads and pathogen shedding potential assessed following surface sampling of the infection site. Mean values ± SEM shown (n=5 per group per time point; 3 technical replicates per fish per time point). **(B)** Effect of temperature on *Aeromonas* growth assessed *in vitro* (n=3 per group per time point). Results analyzed by an ordinary two-way ANOVA using Tukey post-hoc test. Differences correspond to statistical significance of *p<*0.05 between 16°C vs 26°C (**●**), 16°C vs 21°C (♦) and 21°C vs 26°C (▪); ‡ denotes significant difference from time 0, *p<*0.05. **(C)** Hematoxylin and eosin were used to stain infected wounds. ImageJ analysis assessed differential cell recruitment (n=3 per group per time point); refer to Suppl. Figure 2 for source data. Results analyzed by an ordinary two-way ANOVA using Tukey post-hoc test. §corresponds to statistical significance of *p<*0.01 when dynamic and static 26°C groups were compared to 16°C static conditions. No significant differences found between dynamic and static 26°C groups at any time point. Following *Aeromonas* challenge, cutaneous leukocytes were isolated from fish held at 16°C static, 26°C static, or fever dynamic thermal conditions. Imaging flow cytometry evaluated leukocyte production of **(D)** ROS via CellROX, and **(E)** nitric oxide production via DAF-FM-DA (n=5 per group per time point). **(F)** qPCR analysis of wound tissue shows the kinetics of iNOS gene expression (n=5 per group per timepoint; 3 technical replicates per fish per time point). **(G)** Representative ImageStream MKII flow cytometer images show positive and negative cells following staining with CellROX and DAF-FM-DA. Results were analyzed using a two-way ANOVA followed by Tukey’s post-hoc test. \**p<*0.05, \*\**p<*0.01, \*\*\**p<*0.001, \*\*\*\**p<*0.0001; ‡ denotes significant difference from time 0, *p<*0.05.

To assess the contributions of fever to immune cell function, we first quantitated their recruitment into the mucosal infection site. Both T_D_ and T_S26_ fish showed equivalent accelerated infiltration kinetics compared to *Aeromonas*-infected fish held at 16°C static thermal conditions (**Figure 4C** and **Figure S2**). To determine if this enhancement was paired with activation of antimicrobial pathogen killing mechanisms, we then examined the production of reactive oxygen species (ROS) by infiltrating leukocytes, as a prominent, effective, and evolutionarily conserved innate defense mechanism. ^39, 40^ As previously shown by us and others, ^33, 34, 40^ leukocytes derived from fish housed under thermally static conditions (T_S16_ fish for this study) display strong capacity for generation of ROS; over 75% of peritoneal leukocytes were positive for ROS production in the *A. veronii in vivo* cutaneous challenge model (**Figure 4D**). Mechanical hyperthermia (T_S26_) yielded a similar response, with prominent ROS production peaking at 36 hpi (**Figure 4D**). Surprisingly, the number and proportion of ROS-producing leukocytes were greatly reduced under host-driven dynamic thermoregulatory conditions (T_D;_ **Figure 4D**) despite the enhanced kinetics in leukocyte recruitment outlined above (**Figure 4C**).

Given the long-established contributions of fever to host survival, ^3, 9, 20^ we hypothesized that fever may promote an innate antimicrobial response that does not include a strong ROS production component. Thus, we also evaluated leukocyte nitric oxide (NO) production, as an alternative evolutionarily conserved innate response to pathogen attack. ^40, 41^ Contrary to results for ROS, we identified greater overall levels as well as accelerated kinetics of NO production under fever conditions (**Figure 4E**). Leukocytes infiltrating the furuncle of T_D_ fish showed significant upregulation, with peak NO production observed at 24 hpi (**Figure 4E**). This was further supported by a marked, earlier upregulation of inducible nitric oxide synthase (iNOS) expression, which is necessary for production of immune NO (**Figure 4F**). ^40, 41^ In sharp contrast, both T_S16_ and T_S26_ fish displayed lower levels of iNOS expression and lower overall capacity to produce NO (**Figure 4E-G**). Thus, fever differentially regulated ROS and NO leukocyte antimicrobial mechanisms in *Aeromonas*-challenged fish.

### Fever promotes earlier resolution of acute inflammation

To date, studies looking at the basis for host survival due to fever have focused on the activation of immune defense mechanisms. The self-resolving nature of our teleost model allowed us to also characterize immunological changes during the transition between induction and resolution phases of acute inflammation. Indeed, comparison of cellular responses following *Aeromonas* infection showed differences in the control of leukocyte recruitment to the infection site. T_D_ and T_S26_ fish reached the peak of infiltration at 24 hpi, subsequently decreasing and nearing basal levels by 48 hpi (**Figure 4C**). This was consistent with faster kinetics of induction and control of local TNFα, IL-1β, and CXCL8 gene expression among these fish (**Figure 5A**). In contrast, T_S16_ fish showed slower kinetics of leukocyte recruitment with a delayed peak at 48 hpi that was further sustained beyond 72 hpi (**Figure 4C**). These fish also showed delayed upregulation of local TNFα, IL-1β, and CXCL8 gene expression; the latter two pro-inflammatory cytokines further displayed markedly higher levels of expression (**Figure 5A**). Finally, we identified earlier and more pronounced expression of the anti-inflammatory cytokine (TGF-β) and the pro-reparative vascular endothelial growth factor (VEGF) among T_D_ and T_S26_ fish (**Figure 5A**).

**Figure 5.**
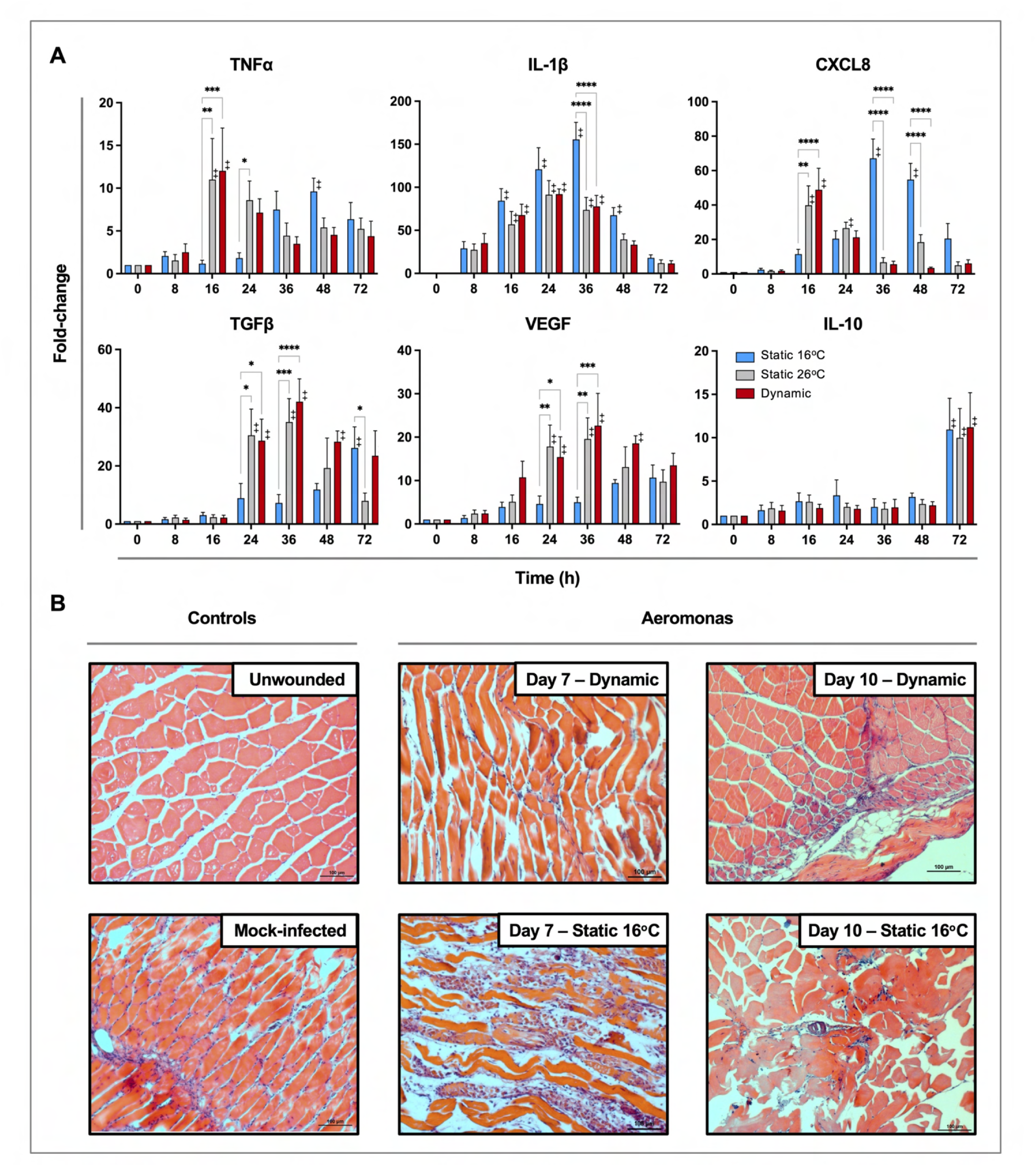
Fever promotes inflammation control and shows early engagement of tissue repair regulators following *Aeromonas* infection. **(A)** Cutaneous wounds were evaluated for expression of pyrogenic cytokines TNF-α and IL-1β, CXCL8 chemokine, and pro-resolution cytokines VEGF, TGF-β and IL-10 via qPCR (n=5 per group per time point; 3 technical replicates per fish per time point). Data was analyzed with an ordinary two-way ANOVA using a Tukey post-hoc test. \**p*<0.05, **p<0.01, \*\*\**p<*0.001, \*\*\*\**p<*0.0001; ‡ denotes significant difference from time 0, *p<*0.05. **(B)** Histopathological assessment highlights inflammation, skin barrier damage and repair at mid and late stages of the infection process. Hematoxylin and eosin stained tissues sectioned from fish inoculated with *Aeromonas* and allowed to exert fever (dynamic) or placed under 16°C static conditions. H&E-stained tissues sectioned from healthy controls (Day 0; non-inoculated) or mock infected fish (Day 0 saline) are provided as controls. n=3 for each group. Scale bar: 100 μm.

Histological examination of experimental wounds further supported marked differences in the inflammation control process of fish maintained in static (T_S16_) and dynamic (T_D_) temperature conditions (**Figure 5B**). Whereas unwounded controls had normal muscle fibers and no signs of inflammation, large infiltrates of granulocytes and macrophages remained in the subdermal muscle tissue of T_S16_ fish 7 days after cutaneous infection with *Aeromonas* (**Figure 5B**). Groups of inflammatory infiltrates were evident in the extracellular space between muscle fibers as well as over damaged muscle fibers. This was in stark contrast to fish allowed to exert fever. Day 7 wounds from these T_D_ fish showed less tissue damage and only traces of the original leukocyte infiltrates remained (**Figure 5B**), thereby resembling those found in the uninfected wounded day 0 control fish (**Figure 5B**). By day 10, infected wounds from fish housed under 16°C static thermal conditions showed prominent necrosis among muscle fibers, edema, and some immune cell infiltrates (**Figure 5B**). In contrast, wounds from fish allowed to exert fever had no necrotic regions and inflammation had largely resolved (**Figure 5B**). Thus, fever promoted an earlier pro-inflammatory period, which was further paired with more efficient resolution of inflammation based on inhibition of leukocyte recruitment, control of pro-inflammatory cytokine expression, induction of pro-resolution genes, and management of collateral tissue damage.

### Fever enhances wound repair

Characterization of pathology at the infection site showed marked differences in the capacity of T_D_ fish to heal *A. veronii*-associated wounds. Similar levels of inflammation were evident in furuncles of T_D,_ T_S26,_ and T_S16_ fish one day after cutaneous infection (**Figure 6**). However, consistent with the enhanced kinetics of leukocyte recruitment (**Figure 4C**), T_D_ and T_S26_ fish showed accelerated kinetics of purulent exudate formation by 2 dpi. Fish exerting dynamic fever subsequently progressed most rapidly, displaying early signs of tissue repair and scale regeneration by 7 dpi, and advanced stages of wound healing by 14 dpi (**Figure 6**). Comparatively, T_S26_ and T_S16_ furuncles did not reach equivalent stages of wound closure (**Figure 6**; green boxes). Thus, fish allowed to exert fever resolved *Aeromonas* infection and repaired the associated skin barrier damage faster than those maintained under mechanical fever-range hyperthermia or static 16°C temperature housing conditions.

**Figure 6.**
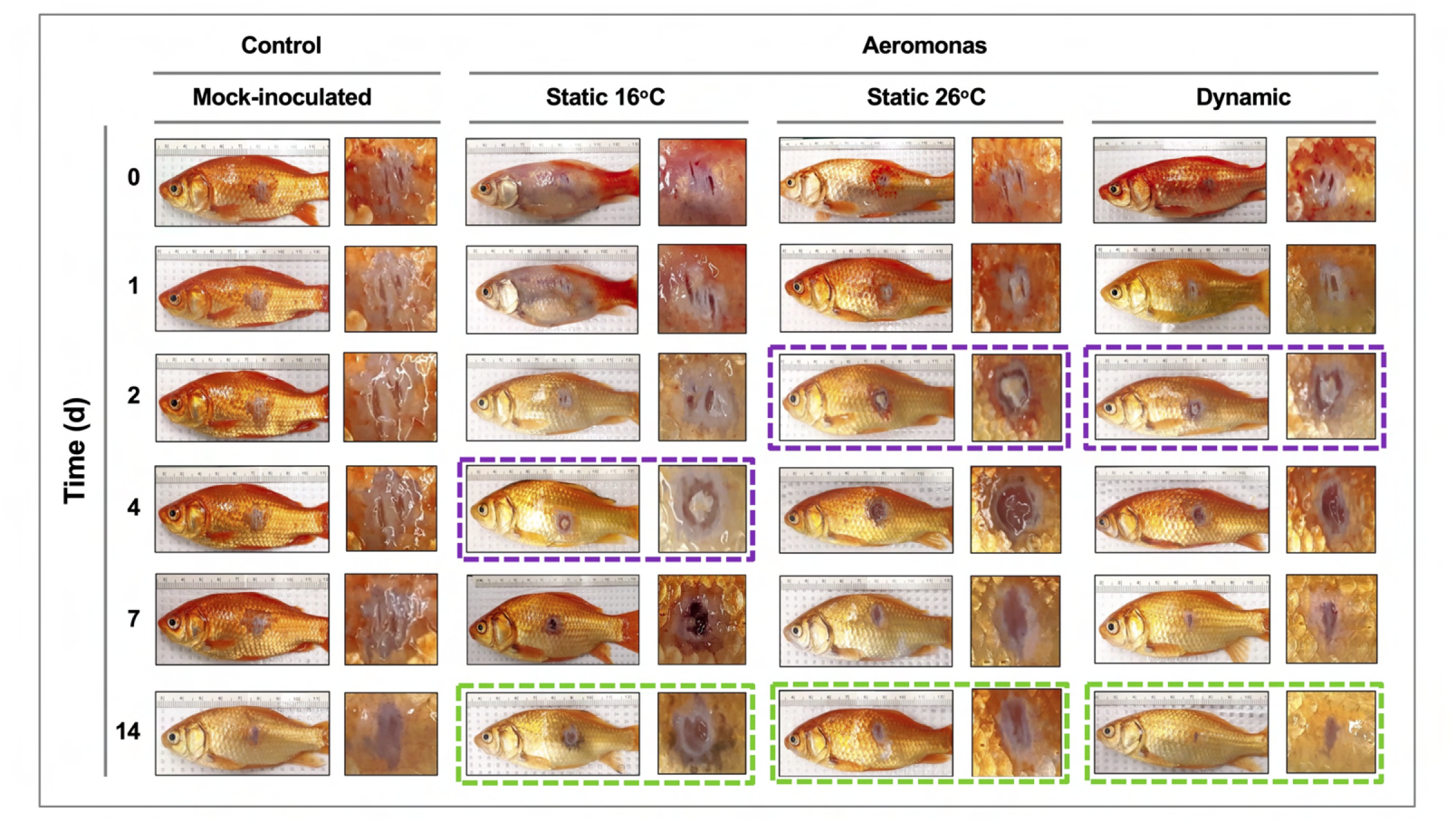
Progression of tissue pathology in *Aeromonas* infected fish. Representative images show focal gross lesions for fish inoculated with *Aeromonas veronii* and placed under 16°C static, 26°C static (mechanical fever-range hyperthermia), or dynamic host-driven fever conditions. Fish mock-infected with saline are provided as controls. Timepoints capture progression from initial infection to advanced stages of wound repair. Purple boxes highlight differential kinetics of purulent exudate formation. Green boxes showcase distinct degrees of wound closure achieved across *Aeromonas*-infected groups by 14 dpi.

Histopathological examination of *Aeromonas*-infected furuncles stained using Masson’s Trichrome stain demonstrated earlier re-establishment of the basal epidermal layer and overlying layers of keratinocytes among T_D_ and T_S26_ experimental groups (Day 3; **Figure 7**). Among T_16_ fish, wound re-epithelialization occurred but was delayed. As per surface pathology described above, tissue repair subsequently advanced faster in wounds derived from T_D_ fish, with *de novo* collagen synthesis becoming evident as early as 4 dpi (**Figure 7**). This further developed into more extensive, organized collagen deposition by 7 dpi (**Figure 7**). In comparison, slower progression was observed among T_S26_ and T_16_ wounds based on the abundance and relative organization of collagen arrangements at the wound site. Regeneration of mucus-secreting cells, consistent with re-establishment of skin barrier functionality, was only observed in the T_D_ experimental group (Day 14; **Figure 7**). Thus, fever promoted greater levels of wound repair and regained original structural features required for restoration of skin barrier functionality after cutaneous infection. Conversely, the absence of fever caused delayed resolution of the inflammatory response, re-epithelialization, and the appearance of extracellular matrix components. Together, our results show that in this cold-blooded vertebrate moderate self-resolving fever offers a natural strategy to harness the body’s intrinsic repair mechanisms to enhance wound healing.

**Figure 7.**
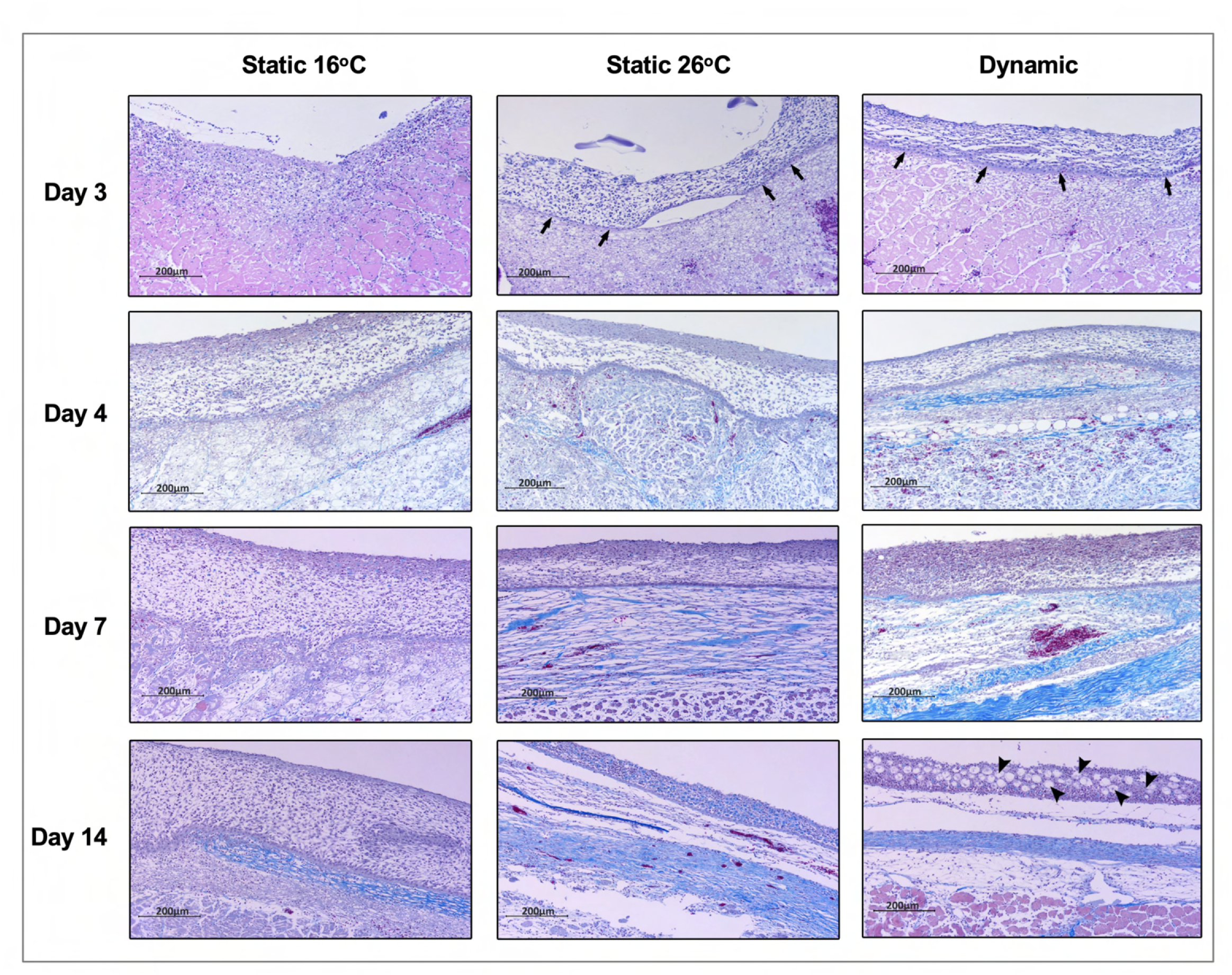
Fever enhances re-epithelialization and collagen deposition. Wound tissue from *Aeromonas* infected fish were collected at the indicated time points, sectioned, and stained with Masson’s Trichrome stain (n=3 per group per time point). Histopathological examination highlights early development of basal layer of epidermis (arrows; Day 3) and overlying layers of keratinocytes among fish in 26°C static and dynamic host-driven fever conditions. Subsequent differential progression of collagen deposition is shown by the increased abundance and organization of blue-stained fibers. Restitution of mucus producing cells in epidermis is highlighted by arrowheads (Day 14). Scale bar: 200 μm.

## DISCUSSION

Induction-centric models cannot explain the continued selection of fever through 550 million years of metazoan evolution, given the predicted high energetic costs of a hypermetabolic state and the implications of inflammation-associated tissue damage. ^14, 24, 27^ A recent emphasis on pathological forms of fever (highest temperatures or sustained hyperthermia; severe disease phenotypes) ^42–44^ further shift our focus away from the more common moderate forms of this natural biological process. ^3, 25^ Thus, for this study we examined fever under an acute inflammatory state that was transient and self-limiting. A eurythermic teleost fish model offered fine control of febrile mechanisms through host-driven dynamic thermoregulation, more closely mimicking natural conditions for heating and cooling. A custom animal enclosure delivered multi-day thermal gradient stability without the use of physical barriers that are known to affect behaviour. ^30, 32^ Together, this combination of animal model, enclosure design, and automated continuous per second tracking of animal locomotory patterns over both induction and resolution phases of the febrile period greatly enhanced analytical robustness and temporal resolution compared to previous studies. Collectively, our data shows that moderate self-resolving fever offers earlier and selective rather than stronger induction of innate antimicrobial programs against infection, and that this results in markedly faster pathogen clearance. We also show that this is paired with rapid subsequent inflammation control and enhanced tissue repair at the wound site. This integrative approach is novel and represents a marked upgrade in refinement compared to popular hypotheses that postulate global induction of immunity, or rely on a shift away from those temperatures preferred by invading pathogens. Our demonstration that this integrative strategy was already well established prior to the development of endothermy further suggests a plausible scenario for the long-standing selection of fever through evolution.

Increases in core body temperature are well established to promote neutrophil accumulation, NADPH oxidase activity, and production rates of toxic superoxide anions. ^45^ Models of fever-range hyperthermia (FRH) have linked these effects to higher serum concentrations of IL-1, TNF, and IL-6, ^46, 47^ G-CSF-driven release of neutrophils from the hematopoietic bone marrow, ^48, 49^ expansion of the circulating neutrophil pool, ^48, 49^ increased endothelial barrier permeability in blood vessels, ^50^ and upregulation of GM-CSF, extracellular HSP70, IL-8 and other CXC chemokines at the local site of infection. ^51, 52^ However, these thermal increases have also been shown to promote collateral tissue injury. ^25, 53, 54^ This continues to be broadly viewed as an unavoidable cost of fever, where induction of immune defenses will undoubtedly drive accompanying inflammation-associated tissue injury. ^51, 54^ Our results offer an alternative explanation, where FRH only partially recapitulates the regulatory capacity of natural fever on acute inflammation. We found that both fever and mechanical fever range-hyperthermia displayed accelerated kinetics of leukocyte recruitment, and earlier control of pro-inflammatory cytokine expression. But differences were observed in the induction of pro-resolution genes, and fever ultimately achieved greater levels of wound repair. We also documented a clear shift from a ROS dominant microbicidal response under euthermic and FRH conditions to one dominated by NO production under fever. NO and downstream reactive nitrogen species (RNS) exert microbicidal or microbiostatic activity against a broad range of bacteria, viruses, yeasts, helminths, and protozoa. ^55^ Yet, at first glance, this inhibition in ROS production seemed contradictory to the role of fever in pathogen resistance. Notably, decreased ROS activity does not necessarily have to result in compromised host defenses. In a murine *Pseudomonas aeruginosa* infection, NO enhanced bacterial clearance via an Atg7-mediated mechanism that also reduced IFN-γ activity, inhibited ROS production, and limited oxidative stress, resulting in decreased lung injury and lower infection-associated mortality. ^56^ A growing number of examples for antagonistic NO modulation of ROS responses during infection have also been described, some of which can be traced as far back as plants. ^39, 55, 57–59^ Our results are consistent with these observations and offer a natural scenario where fever drives a shift in the NO-ROS balance that maintains competencies in microbial clearance to subvert a live infection, while also contributing to inflammation control and the re-establishment of a functional mucosal barrier.

Recent years have seen renewed interest in the links between thermoregulation and host defense, which now also extend to cooler temperatures. ^13, 14, 43, 60, 61^ This has led to a perceived functional dichotomy between fever and hypothermia. Although both reflect an animal’s capacity to take advantage of thermoregulation to maintain fitness upon infection, fever drives elimination of invading microorganisms through microbicidal disease resistance while hypothermia promotes tolerance to foster energy conservation and management of collateral inflammation-associated tissue damage. ^13, 14^ Our results blur the line in this perceived dichotomy. We identified clear fever-embedded mechanisms that contributed to the maintenance of tissue integrity and would allow for energy conservation. These contributions did not appear in response to damage, but instead were prominent features through initial induction and subsequent resolution phases of acute inflammation. Early on, fever promoted disease resistance via enhanced kinetics of leukocyte recruitment rather than by augmenting the magnitude or duration of immune activation. Engagement of cytoprotective gene programs in the CNS occurred within hours of the initial immune challenge. Selective rather than global upregulation of immunity further decreased the potential for collateral damage often attributed to fever. Notably, microbicidal efficacy was still superior under fever, with *Aeromonas* clearance achieved markedly faster than under thermally restricted basal conditions (7 versus 14 days). Our results then showed greater efficiency in inflammation control. This was paired with novel contributions that actively promoted tissue repair rather than resilience to inflammatory damage. There was no induction of a hypothermic state at any point during our observation periods; instead, high-resolution positional tracking only showed a discrete, self-resolving fever response. Thus, fever actively engages mechanisms that enhance protection, while also limiting pathology, controlling inflammation, and promoting tissue repair. Importantly, our findings do not dispute previously described competition mechanisms between immunity and other maintenance programs that direct a transition towards tolerance under high pathogen loads. ^43, 62^ Instead, they simply highlight the persistent and long-standing considerations placed on energy allocation and tissue integrity by an animal host at all stages of infection.

To conclude, our results reveal novel features of fever, and demonstrate that it is an integrative host response to infection that regulates both induction and resolution phases of acute inflammation. The downstream implications of our findings are far-reaching. Among others, enhanced re-establishment of barrier integrity at tissue sites such as the skin are predicted to reduce the potential for secondary infections, and curtail the physiological stress stemming from a need for extended management of wounded tissue. There are also inferences at the population level, where marked enhancements observed in pathogen clearance are likely to translate into lower rates of transmission across a naive population, offering novel opportunities for the management of infectious disease. More studies will be required to test these possibilities, but this is critical as we look to better understand the long-standing role of fever in the modulation of acute inflammation, and the repercussions of inhibiting moderate fever in veterinary and human medicine.

## MATERIALS AND METHODS

### Ethics statement

All animals were maintained according to the guidelines of the Canadian Council on Animal Care. All protocols were approved by the University of Alberta Animal Care and Use Committee (ACUC-Biosciences protocols 706 and 355303). Fish were terminated via cervical dislocations using approved procedures. All efforts were made to minimize animal stress and to ensure that termination procedures were efficiently performed.

### Animals

Goldfish (*Carassius auratus auratus*), 10-15 cm in length, mix-sex, were purchased from Mt. Parnell Fisheries (Mercersburg, PA) and imported to Canada via Aquatic Imports (Calgary, Canada). They were held in a flow through system with simulated natural photoperiod (12 hours of light: 12 hours of dark) in the Aquatics Facility, Department of Biological Sciences, University of Alberta. Water quality parameters were sustained at 5.5–6.5 PPM dissolved oxygen and pH 7.2–8.0. Fish were fed with floating pellets once daily.

### Design and validation of the annular temperature preference apparatus

The ATPT was constructed from customized precision cut acrylic sheets molded and sealed into three concentric rings: an outer-most inflow ring separated into eight equal segments around the periphery, a middle continuous swim chamber containing no physical barriers, and an inner circle used to control depth. An additional inner compartment contained outflow and drainage. Small equidistantly drilled pores placed high on the outer inflow chambers and low on the inner outflow chamber allowed for water flow from the periphery to the center of the apparatus. This ring-shaped swim chamber maintained constant water depth, current and perceived cover throughout, which are factors known to impact behaviour of aquatic animals. Eight distinct thermal zones were maintained by fluid dynamics. Temperatures of these zones were monitored on a per second basis over 14 days using a HOBOware U30 data-logger with 12-bit temperature sensors (Onset Computer Corporation, Bourne, MA).

### Quantification of animal behaviours

Fish behaviour was recorded by a centrally placed overhead infrared camera (Panasonic CCTV colour Camera, WV-CP620 with a 2X Lens variable focal WV-LZ61/2S) and lighting system. This allowed continuous digital video recording of fish movement through simulated day and night cycles. Videos were analyzed using Ethovision XT, Version 11 (Noldus, Wageningen, Netherlands) to quantify behaviours by automated animal tracking. Dynamic subtraction was used to target and track coordinates of each animal within the ATPT on a per second basis, and finished tracks were manually verified. The field of view was separated into eight zones corresponding to 16°C, 19°C (L, R), 21°C (L, R), 23°C (L, R), or 26°C (**Figure 1**). This was used to calculate fish preference for each thermal zone, velocity of movement, and migration between thermal zones. Data was compiled and used to calculate mean hourly temperature preference, velocity, and number of transitions across thermal zones.

### *In vivo Aeromonas* infection model and quantification of pathogen loads on skin wounds

*Aeromonas veronii* biovar sobria (NCBI Taxonomy ID: 114517) was previously isolated by our lab from goldfish cutaneous lesions. ^33^ For culture preparation, bacteria were inoculated into a 5 mL sterile trypticase soy media (BD Biosciences, Franklin Lakes, NJ) and cultured at room temperature on a shaker overnight. Fish were anesthetised in a tricaine methanesulfonate (TMS) solution (02168510; Syndel, WA), a 4 × 4 patch of scales was removed, and minor abrasions made on the skin. The cutaneous wound was then inoculated with 10 µL of *A. veronii* log-phase culture broth (4.1 × 10^8^ CFU/mL) before returning the fish to water. This inoculation dose was previously determined to promote a self-resolving acute inflammatory process where initial induction and subsequent resolution phases could be examined. ^33^ Infected fish were assigned randomly to different temperature categories, and they were also randomly selected at indicated time points for our experiments. Subsequent assessment of *A. veronii* numbers at the furuncle surface was used as a measure of pathogen load and shedding potential; surface bacteria were collected and plated onto 2x replicate tryptic soy agar plates. CFUs were quantified after 24 h incubation at room temperature. Ketorolac (02162644; ATNAHS, Basildon, UK) administered intraperitoneally at 0.5 mg/kg of body weight was used in select experiments.

### Gene expression

Samples were collected and immediately frozen in liquid nitrogen and stored at −80 °C until use. Total RNA was extracted using TRIzol (15596026; Thermo Fisher Scientific, Waltham, MA) following the manufacturer’s specifications. RNA concentration and quality were evaluated using a Nanodrop ND-1000 (Thermo Fisher Scientific, Waltham, MA) and Bioanalyser-2100 equipped with an RNA 6000 Nano Kit (5067-1511; Agilent Technologies, Santa Clara, CA). cDNA was synthesized using iScript Kit (1708891; BioRad, Mississauga, Canada) according to the manufacturer’s specifications. qPCR was performed using the QuantStudio 6 Flex Real-Time PCR System (Applied Biosystems, Waltham, MA) where RQ values were normalized against gene expression on day 0 for each replicate time course and β-actin was used as a reference gene. Primers used are listed in **Table S1**. Relative quantification was performed according to the 2*^−ΔΔCt^* method.

### ROS production

Leukocyte isolations and ROS production evaluations were performed as previously described. ^34, 63^ After isolations, 500 µL of cell suspensions were incubated with 0.5 µL of CellROX™ Deep Red Reagent (C10491; Thermo Fisher Scientific, Waltham, MA) in the dark for 30 min to allow for cellular uptake. Cells were washed twice with 1 × PBS^−/−^ and fixed with 1% formaldehyde (47608; Sigma Aldrich, St. Louis, MO). Samples were centrifuged at 350 × g for 5 min at 4°C. Data was acquired using an ImageStream Mk II Imaging Flow Cytometer (Amnis, Seattle, WA), and analyzed via IDEAS Image Data Exploration and Analysis Software (Amnis, Seattle, WA). Cells were gated based on the normalized frequency of a fluorescent minus one sample.

### NO production

The production of NO was evaluated using 4-amino-5-methylamino-2’,7’-difluorofluorescein diacetate (DAF-FM DA)(D23844; Invitrogen, Waltham, MA). Following leukocyte isolation, 500 µL of cell suspensions were incubated with DAF-FM DA at a concentration of 1 µM for 30 min in the dark. Cells were washed twice with 1 × PBS^−/−^ and fixed with 1% formaldehyde (47608; Sigma Aldrich, St. Louis, MO). Samples were centrifuged at 350 × g for 5 min at 4°C. Data was acquired using an ImageStream Mk II Imaging Flow Cytometer (Amnis, Seattle, WA), and analyzed via IDEAS Image Data Exploration and Analysis Software (Amnis, Seattle, WA). Cells were gated based on the normalized frequency of a fluorescent minus one sample.

### Prostaglandin E2 in goldfish plasma

Following cutaneous infection with *A. veronii*, fish were placed in either 16°C static, 26°C static, or dynamic fever thermal conditions. At 0, 12, 24, 48 and 72 h post-infection, heparinized blood was collected from 6 individuals for each thermal condition and centrifuged at 2000 × g for 10 min at 4°C. The resulting plasma supernatant was collected, aliquoted, and stored at −80°C until use. Using a Prostaglandin E2 ELISA Kit (514010; Cayman Chemical, Ann Arbor, MI), plasma samples were diluted 1:30 and PGE_2_ protein concentrations were determined following the manufacturer protocol. Plates were read using the SpectraMax M2e plate reader (Molecular Devices, San Jose, CA) at 405 nm. Analysis and quantification of PGE_2_ protein production was completed as per manufacturer specifications (Cayman Chemical, Ann Arbor, MI).

### Histopathological analysis

Wound tissues were collected and fixed in 10% neutral-buffered formalin (SF98-4; Thermo Fisher Scientific, Waltham, MA)). After processing tissues overnight in a series of ethanol, toluene, and wax washes using a Leica TP1020 benchtop tissue processor (Leica Biosystems, Concord, Canada), they were paraffin-embedded and sectioned (7 µm thickness) on slides using a Leica RM2125 RTS microtome (Leica Biosystems, Concord, Canada). Slides were deparaffinized and washed using two rounds of toluene (T324-1; Thermo Fisher Scientific, Waltham, MA) (5 min each) followed by rounds of 100%, 90%, 70%, and 50% ethanol (2 min each). For Hematoxylin & Eosin (H&E) staining, slides were placed in Surgipath Hematoxylin Gill III (3801542; Leica Biosystems, Concord, Canada) for 2 min, washed with running cold tap water for 15 min followed by 70% ethanol for 2 min, and then in Surgipath Eosin solution (3801602; Leica Biosystems, Concord, Canada) for 30 sec. For Masson’s Trichrome stain, slides were placed in hematoxylin Gill III for 1 min and washed with running cold tap water for 15 min. Slides were stained with ponceau-fuchsin (AC400211000; Thermo Fisher Scientific, Waltham, MA) for 2 min, rinsed in distilled water, differentiated in mordant in 1% phosphomolybdic acid (19400; Electron Microscope Sciences, Hatfield, PA) for 5 min. Slides were then stained with Aniline Blue solution (A967-25; Thermo Fisher Scientific, Waltham, MA) for 3 min and incubated in 1% phosphomolybdic acid then acetic acid solution (A38C-212; Thermo Fisher Scientific, Waltham, MA) for 5 min and 3 min, respectively. Lastly, for both H&E and Masson’s Trichrome, slides were dehydrated in series of alcohol, cleared in toluene, mounted with DPX Mountant (50-980-370; Thermo Fisher Scientific, Waltham, MA). Images were obtained using an AxioScope A1 microscope (Zeiss, Oberkochen, Germany).

### Statistics

Data was statistically analyzed and graphed using GraphPad v9.3.1 (San Diego, CA). Fish numbers used in each experiment were determined based on the minimal sample size (n) required to detect statistical significance, while considering animal ethics and care guidelines under the Canadian Council on Animal Care. Welch’s t-test was used to determine if the means of two groups were significantly different. Data of both groups had a normally distributed population, but it was not assumed that they had the same variance. One-way ANOVA was used to compare the variance in the means of three or more categorical independent groups, considering a single independent factor or variable in the analysis. Data had a normally distributed population and each sample was drawn independently of the other samples. Additionally, the dependent variable was continuous. Two-way ANOVA was utilized for comparing the effect of two independent categorical factors on a dependent variable such as fold-change. The dependent variable was continuous and each sample was drawn independently of the other samples. Šídák’s multiple comparisons test was used in select cases when a set of means were selected to compare across two groups and each comparison was assumed to be independent of the others. Mean values and correlations for behavioural data were calculated in Excel (Microsoft, Redmond, WA). R (version 3.3, The R Foundation for Statistical Computing, Vienna, Austria) was used to calculate multivariate statistics including principal component analysis (standard R package) and permutational multivariate analysis of variance using distance matrices (‘vegan’ community ecology package). R code is included as part of the supplementary material.

## ABBREVIATIONS

ATPT: annular temperature preference tank
CNS: central nervous system
COX: cyclooxygenase
DAMPs: damage-associated molecular patterns
DPI: days post-infection
FRH: fever-range hyperthermia
G-CSF: granulocyte colony-stimulating factor
GM-CSF: granulocyte-macrophage colony-stimulating factor
HPI: hours post-infection
HSF1: heat shock factor 1
HSP: heat shock protein
IFN: interferon
IL: interleukin
iNOS: inducible nitric oxide synthase
NSAIDs: Non-steroidal anti-inflammatory drugs
NO: nitric oxide
PAMPs: pathogen associated-molecular patterns
PGE2: prostaglandin E2
PRRs: pattern recognition receptors
RNS: reactive nitrogen species
ROS: reactive oxygen species
T_D_: dynamic thermoregulation (fever)
T_S16_: static thermal conditions at 16°C
T_S26_: static thermal conditions at 26°C
TGF-β: transforming growth factor-β
TNF: tumor necrosis factor
V: velocity
VEGF: vascular endothelial growth factor
ZT: zone transitions

## ACKNOWLEDGEMENTS

We thank M. Reichert, C. Gerla, J. Edgington, M. Axelsson, and J. Johnston for help in the construction of the custom ATPT, the Science Animal Support Services for maintaining experimental fish, and A. Shostak and Z. Song for input on statistics and multivariate analyses. This work was supported by a Natural Sciences and Engineering Council of Canada grant to DRB (RGPIN-2018-05768). AMS and MEW were supported by a Graduate Teaching Assistantship from the Department of Biological Sciences at the University of Alberta. DT was supported by a CONICYT-Chile postdoctoral fellowship (Becas Chile N° 74170029). SLS was supported by a Natural Sciences and Engineering Council of Canada post-doctoral fellowship. The funders had no role in study design, data collection and analysis, decision to publish, or preparation of the manuscript.

## DATA AVAILABILITY

All data and materials generated or analyzed during this study are included in this manuscript or provided as supplementary material.

---

**Supplementary figure 1:**
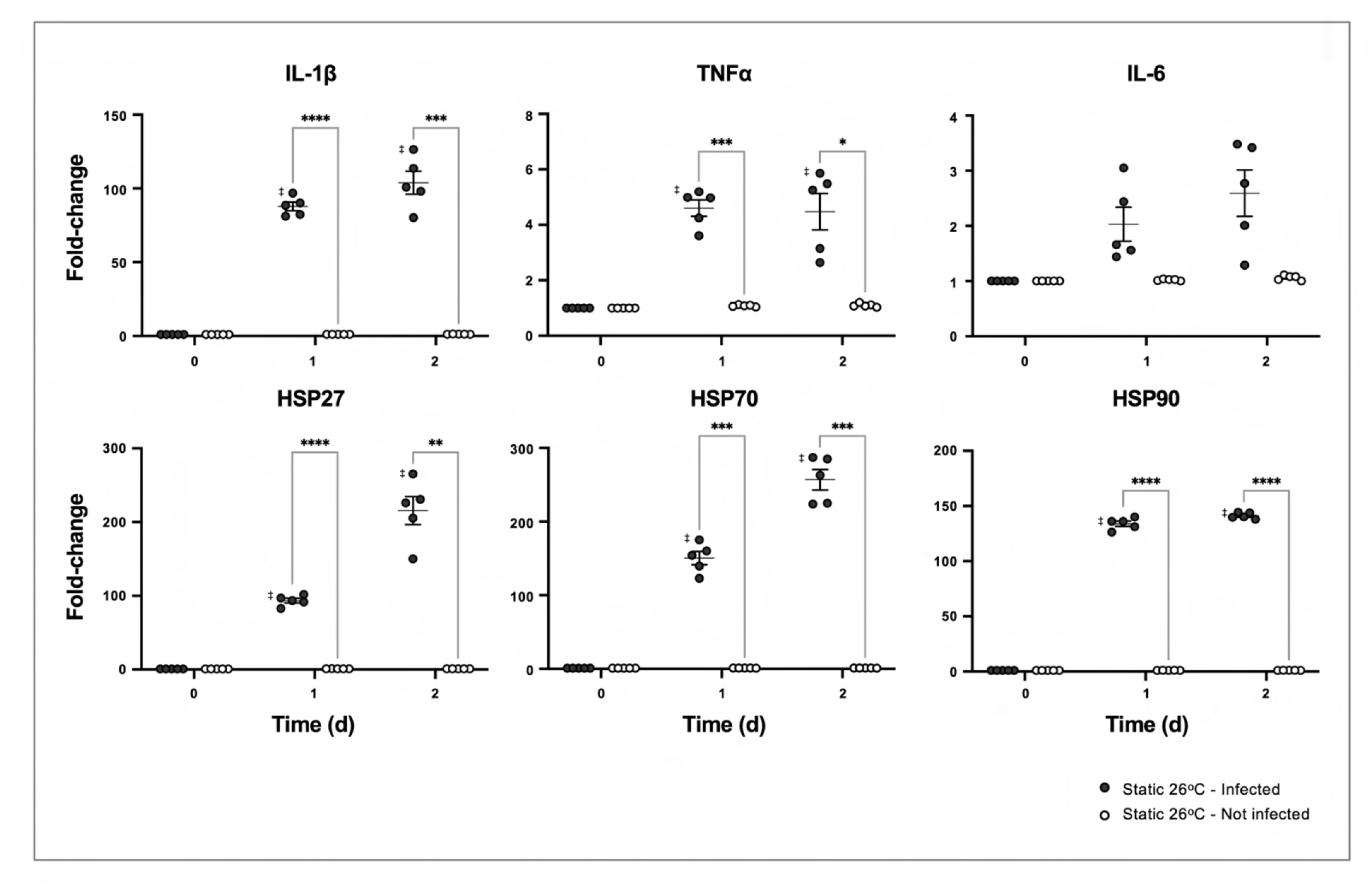
Thermal increases alone are not sufficient to engage CNS and stimulate immune regulatory and cytoprotective genes. *Aeromonas*- or mock-infected fish were placed in 26°C static thermal conditions following inoculation. Quantitative real-time PCR evaluated gene expression in the hypothalamus following infection (n=5 per group per time point; 3 technical replicates per fish per time point). Symbols correspond to individual samples and lines represent the mean ±SEM. Results were analyzed by an ordinary two-way ANOVA and Šídák’s multiple comparisons test. \**p<*0.05, \*\**p<*0.01, \*\*\**p<*0.001, \*\*\*\**p<*0.0001; ‡ denotes significant difference from time 0, *p<*0.05.

**Supplementary figure 2:**
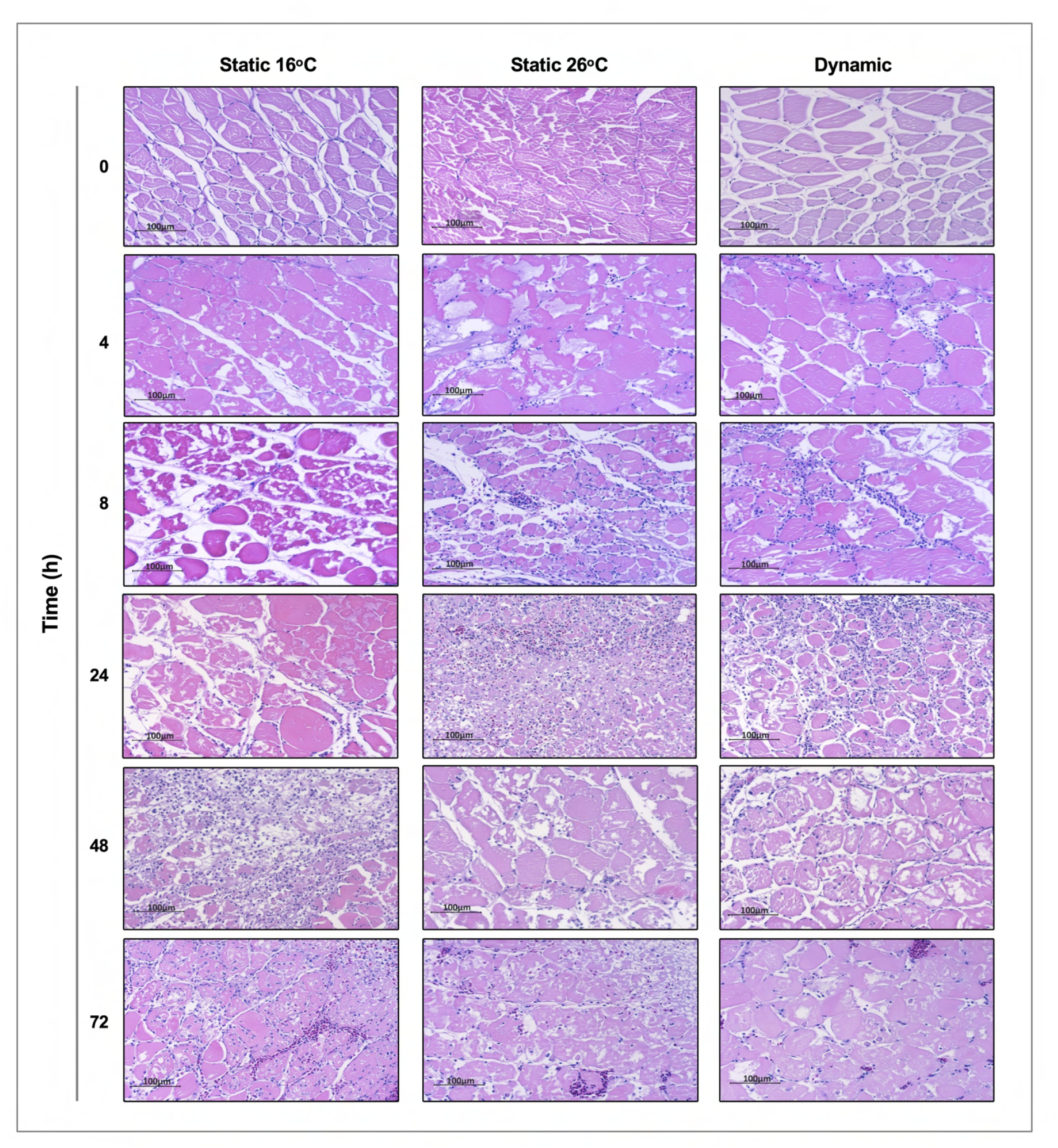
Thermal promotion of leukocyte recruitment to skin wounds in *Aeromonas* infected fish. Hematoxylin and eosin were used to stain tissues sectioned from the skin infection site after housing of fish at 16°C static, 26_o_C static, or dynamic host-driven fever conditions. Timepoints capture initial 0-72 h of acute inflammation (n=3 per group per time point). Scale bar: 100 μm.

**Supplementary table 1: Primer sequences for quantitative PCR**

